# Evaluation of quantum dot conjugated antibodies for immunofluorescent labelling of cellular targets

**DOI:** 10.1101/090357

**Authors:** Jennifer E. Francis, David Mason, Raphaël Lévy

## Abstract

Semiconductor quantum dots (Qdots) have been utilised as probes in fluorescent microscopy and provide an alternative to fluorescent dyes and fluorescent proteins, due to their brightness, photostability, and the possibility to excite different Qdots with a single wavelength. In spite of these attractive properties, their take up by biologists has been somewhat limited and only a few Qdot conjugates are commercially available for the labelling of cellular targets. Although, many protocols have been reported for the specific labelling of proteins with Qdots, the majority of these relied on Qdot-conjugated antibodies synthesised specifically by the authors and therefore not broadly available, which limits the scope of applications and complicates replication. Here, the specificity of a commercially available Qdot conjugated secondary antibody (Qdot-Ab), for different antigens, was tested. Antigens were labelled simultaneously with a fluorescent dye coupled to a secondary antibody (Dye-Ab) and the Qdot-Ab. Although, the Dye-Ab labelled all of the intended target proteins, the Qdot-Ab only bound to some of the protein targets in the cytosol and could not reach the nucleus even after extensive cell permeabilisation.

## Introduction

Quantum dots (Qdots), nanometer-sized semiconductor nanocrystals, that typically consist of a metallic core of cadmium selenium (CdSe) and an inorganic zinc sulfide (ZnS) shell, have been applied as fluorescent probes for the labelling of biological structures [1, 2]. To make Qdots water-soluble and thus suitable for biological applications, their surface is either modified by coating with hydrophilic ligands, such as poly(ethylene glycol) (PEG) [3, 4], or encapsulated in amphiphilic polymers [5]. Antibodies that recognise specific biological targets can then be conjugated to these Qdots for use in immunofluorescence. Conventional immunocytochemistry (ICC) protocols involve the chemical fixation of cells, followed by permeabilisation with a detergent. This creates pores in the cell membrane, allowing primary and secondary antibodies to gain access to the protein of interest.

Qdots are an attractive alternative to traditional fluorescent dyes for ICC, because they are much brighter and more photostable [6, 7]. In contrast to fluorescent dyes, Qdots can be excited with a wide range of wavelengths and have narrow emission spectra, which is advantageous for multiplex imaging [8, 9]. The emission maxima of Qdots are dependent on their size, with the emission peak for large Qdots being in the red end of the spectra and smaller Qdots in the blue region [1]. Qdots are also an ideal probe choice for super-resolution techniques that require stochastic optical fluctuation, as they have a well characterised blinking between fluorescent and non-fluorescent states [10, 11].

Despite these favourable characteristics, once conjugated to antibodies, the overall hydrodynamic radius of a Qdot (15-20 nm) is much larger than that of a fluorescent dye [12-14]. As a result, one large Qdot may host many antibodies, whereas many fluorescent dye molecules can be coupled to a single antibody [15]. Furthermore, the overall size of commercially available Qdots is further enlarged, by the addition of protective layers to maintain stability and shelf life [13, 16]. Qdot conjugated antibodies (Qdot-Abs) are therefore unlikely to replace fluorescent dyes in ICC, due to their inferior penetration capability [17, 18]. Despite the assumption that manufactured Qdots are quality controlled, considerable batch-to-batch variability, means that each time a new lot is purchased, the labelling conditions need to be optimised [19]. The use of commercial Qdot-Abs is also expensive, as they are to be used at a much higher concentration than fluorescent dyes, in the order of 20 nM [16, 20, 21]. Despite having been around for several decades, Qdots are rarely used in routine ICC [1].

There are some notable examples of Qdot-Abs in the published literature, to label glycine receptors [22], glial fibrillary acidic proteins (GFAPs) [12], mortalin [23], erythrocytes [24], GRP78 protein [25], caveolin-1 [26], golgi [20], and nuclear HER2 targets [6]. The majority of this labelling however, was done with in-house synthesised Qdots rather than commercially available Qdots [27]. In this way, the synthesis can be controlled and the size of the Qdot-Abs kept to a minimum. However, it has been noted that specific labelling of nuclear and some cytoplasmic structures with Qdot-Abs is not always reproducible [1, 6, 13, 20, 28]. Another concern amongst users of commercial Qdot-Abs is non-specific labelling and the formation of aggregates, which may introduce artifacts and lead to misinterpretation of false positive results [12, 13].

Here, we focus on the specificity of commercially available Qdot 625 (emission maxima of 625 nm) conjugated antibodies (Thermo Fisher Scientific, UK) in fixed cells (Scheme 1). Different protein targets were labelled simultaneously with both a secondary antibody conjugated to a fluorescent dye (Dye-Ab) and a Qdot 625 conjugated secondary antibody (Qdot-Ab). Anti-GFP Qdot 625 conjugate (Qdot-GFP) and Qdot 625 conjugated to streptavidin (Qdot-Streptavidin) were also evaluated. We found that while the prototypical target of Qdot-Abs: tubulin, could be easily labelled, several other protein targets including nuclear proteins and components of large cytosolic protein complexes could not be labelled with Qdot-Abs. We posit that this may be due to steric hindrance associated with the size of the Qdot-Abs.

**Scheme 1.**
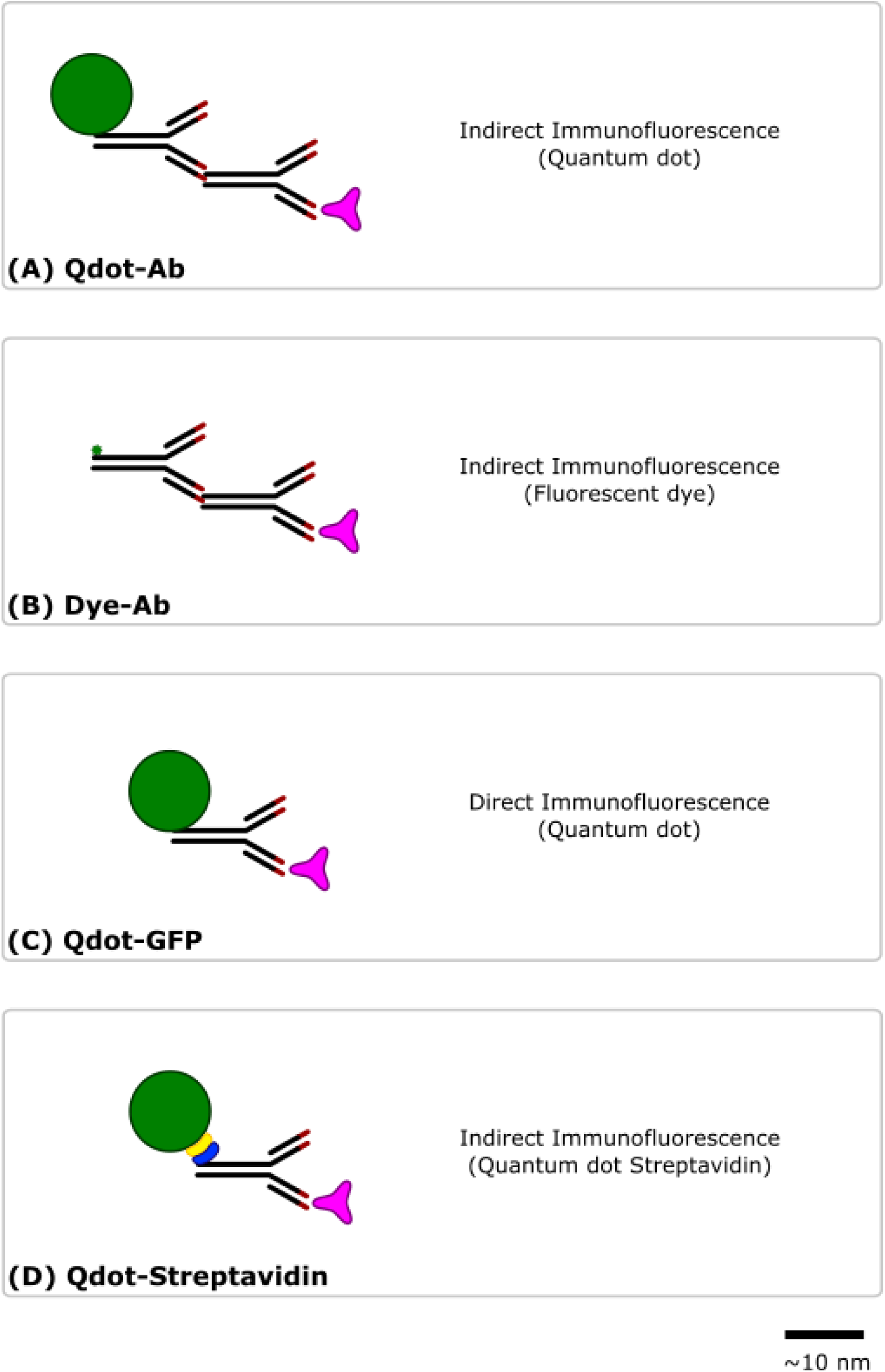
Strategies for immunofluorescence labelling. A fluorescent label (green) is conjugated to a secondary or primary antibody (black), containing an antigen-binding site (red), which recognises and binds to a specific antigen (pink). Immunofluorescence labelling was either indirect with a primary antibody and a Qdot 625 (A) or fluorescent dye (Alexa Fluor 488 or Cyanine 3) (B) conjugated to a secondary antibody, or direct with an anti-GFP primary antibody conjugated to Qdot 625 (Qdot-GFP) (C). An alternative Qdot 625 was tried for indirect immunofluorescence labelling using a Qdot 625 streptavidin (yellow) conjugate (Qdot 625 streptavidin) and biontinylated (blue) primary antibody. Scale bar is 10 nm.

## Results and Discussion

*Labelling of extracellular antigens:* To investigate the specificity of commercial Qdot-Abs for intracellular targets, different types of proteins were stained simultaneously with conventional Dye-Abs and Qdot-Abs and imaged using an epifluorescence microscope. To eliminate the possibility of competition between the Dye-Abs and Qdot-Abs, for the antigen binding sites, which could affect the labelling, samples were also prepared separately with either a Dye-Ab or Qdot-Ab. A Qdot 625 concentration of 20 nM was used, as it has been shown that a high concentration of Qdot-Abs improves specific labelling and signal-to-noise ratio [21]. Initially, to assess the labelling efficiency of Qdot-Abs, the extracellular matrix (ECM) protein fibronectin was dual labelled with Qdot 625 and Dye-Ab (Alexa Fluor 488). Fibronectin is abundant at the cell surface and therefore, would not be expected to display any artifacts associated with accessibility. Indeed, in this case similar labelling was achieved with Qdot 625 and Alexa Fluor 488 (Figure 1).

**Figure 1.**
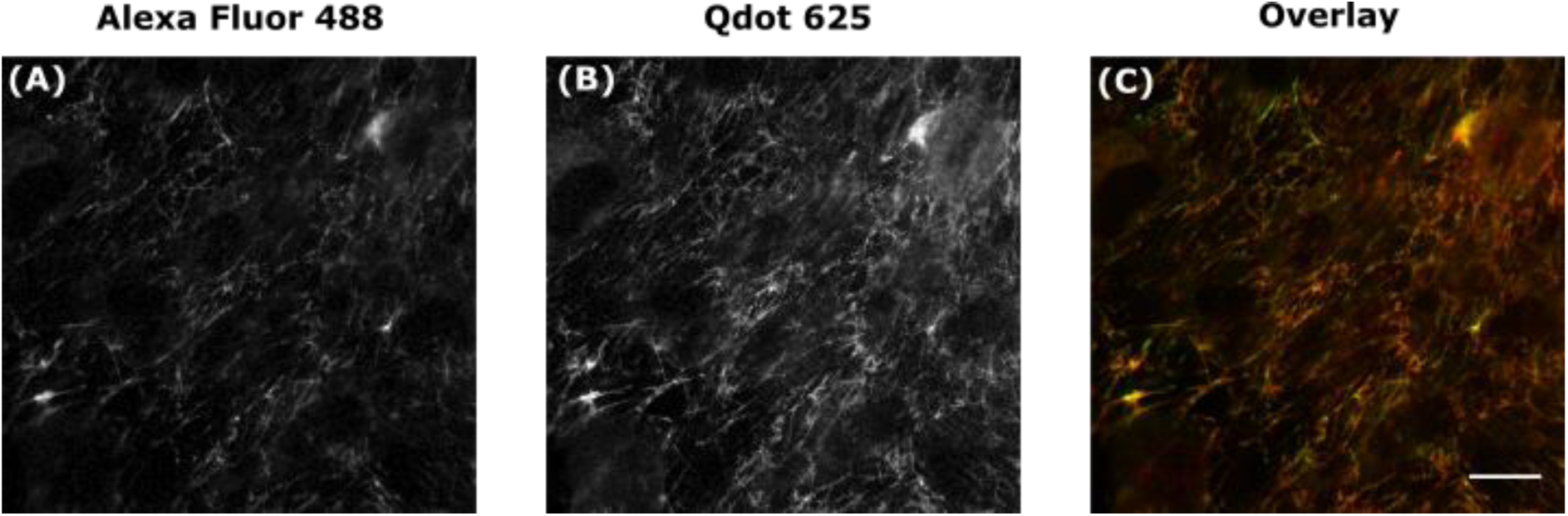
Specific labelling of fibronectin with Qdots. Fixed *rat mammary (Rama) 27 fibroblasts* were dual labelled with green Alexa Fluor 488 (A) and a red Qdot 625 (B) to produce an overlaid wide-field image (C). Scale bar is 20 μm.

*Labelling of cytosolic structures:* The vast majority of studies using Qdots show tubulin staining [6, 12, 16, 18, 20], therefore we sought to label this abundant cytosolic protein in order to have a positive control for our labelling protocol. After incubation with an anti-tubulin primary antibody and a Qdot-Ab, we once again found labelling which was comparable to samples (simultaneously or separately) labelled with Alexa Fluor 488 (Figure 2).

**Figure 2.**
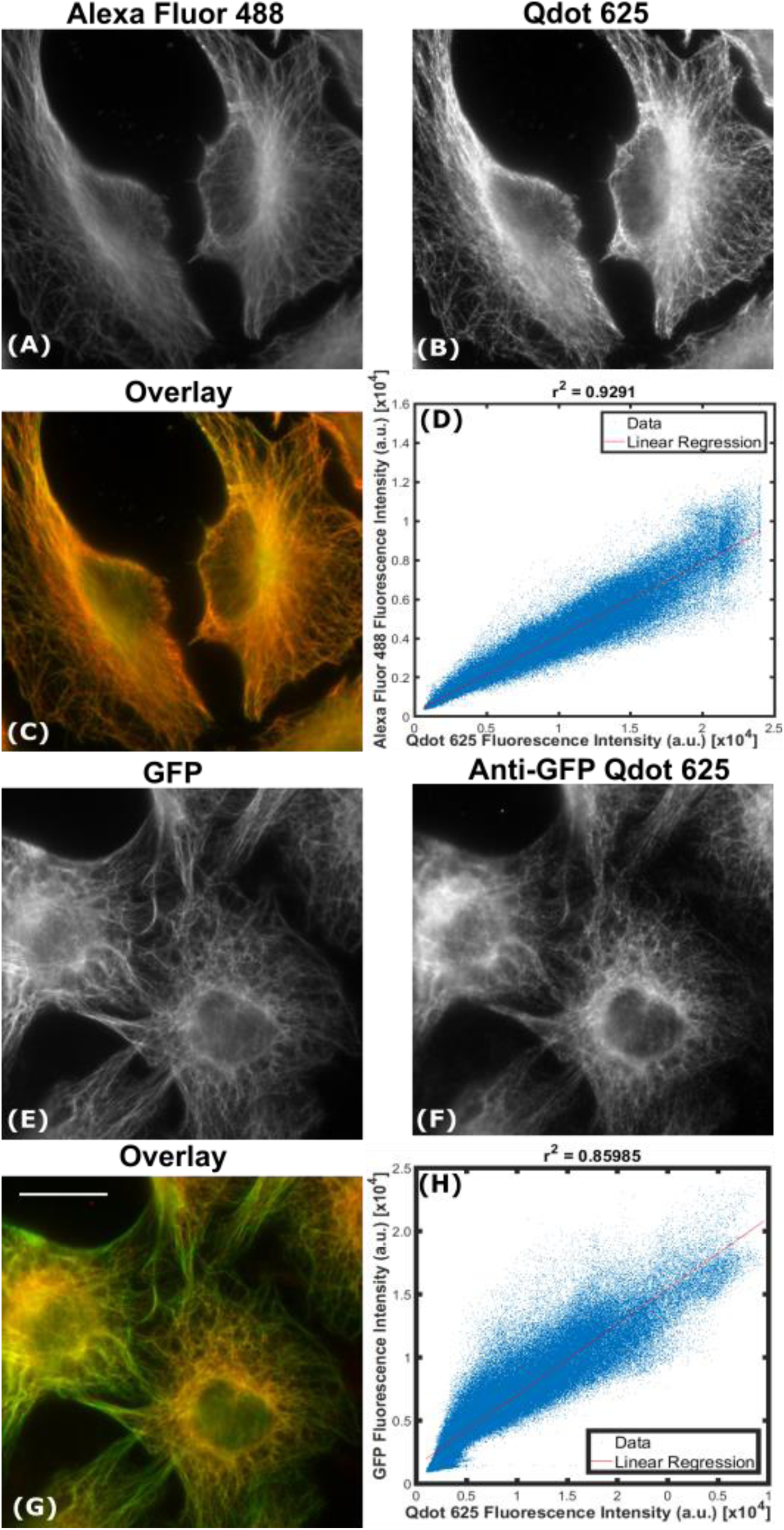
Specific labelling of tubulin with Qdots. Methanol-fixed HeLa cells were labelled indirectly with a primary anti-tubulin antibody, green Alexa Fluor 488 (A), and red Qdot 625 (B), to produce an overlaid wide-field image (C). As a measure of co-localisation between Alexa Fluor 488 and Qdot 625, fluorescence intensities of the overlaid wide-field image (C) were analysed using a custom-written Matlab code to produce a correlation scatter plot (D). Pearson’s correlation coefficient (r^2^) of 0.93 indicates very high correlation. The average Manders coefficient of Alexa Fluor 488 overlapping with Qdot 625 (M1) was 0.99 (SD=0.007, N=3) and the average Manders coefficient of Qdot 625 overlapping with Alexa Fluor 488 (M2) was 0.99 (SD=0.005, N=3, as determined using JACoP.Paraformaldehyde fixed TC7 cells, expressing tubulin-GFP (E), were also labelled directly with an anti-GFP Qdot 625 conjugate (F) to produce an overlaid wide-field image (G,) and a corresponding correlation scatter plot (H); with a Pearson’s correlation coefficient (r^2^) of 0.86. Scale bar is 20 μm.

To investigate whether the same structure (microtubules) could be labelled with a different antigen, direct ICC was performed using a Qdot 625 conjugated directly to an anti-GFP primary antibody (see Methods for details). Cells expressing tubulin-GFP were labelled with the anti-GFP Qdot 625 conjugate (Qdot-GFP) and gave similar results to those imaged with indirect ICC (Figure 2). By inspection it was clear, that β-tubulin had been labelled with Qdot 625 (Figure 2C), however, to compare the two labelling techniques, an intensity correlation plot was produced (Figure 2D). A Pearson’s correlation coefficient (r^2^) value of 0.93 suggested that there was an almost perfect correlation between Qdot 625 and Alexa Fluor 488 labelled microtubules. In addition, the degree of co-localisation was measured using the Manders coefficient [29]. The average Manders coefficient of Alexa Fluor 488 overlapping with Qdot 625 (M1) was 0.99 (SD=0.007, N=3) and Qdot 625 overlapping with Alexa Fluor 488 (M2) was 0.99 (SD=0.005, N=3), confirming that the correlation between Qdot 625 and Alexa Fluor 488 was very good.

*Labelling intracellular complexes*: Both fibronectin and β-tubulin are highly abundant proteins. Furthermore, their antigens are relatively accessible. We therefore looked to antigens present in more complex intracellular structures, including the focal adhesion protein talin and nuclear splicer marker SC35. Talin exists in a dynamic equilibrium with both a bound pool (forming focal adhesions) and a cytosolic pool. Using the same primary antibody, both Alexa Fluor 488 and Qdot 625 labelled the cytosolic pool of talin (Figure 3). Interestingly however, the bound pool forming focal adhesions were only visible when labelled with Alexa Fluor 488, and not Qdot 625 (Figure 3).

**Figure 3.**
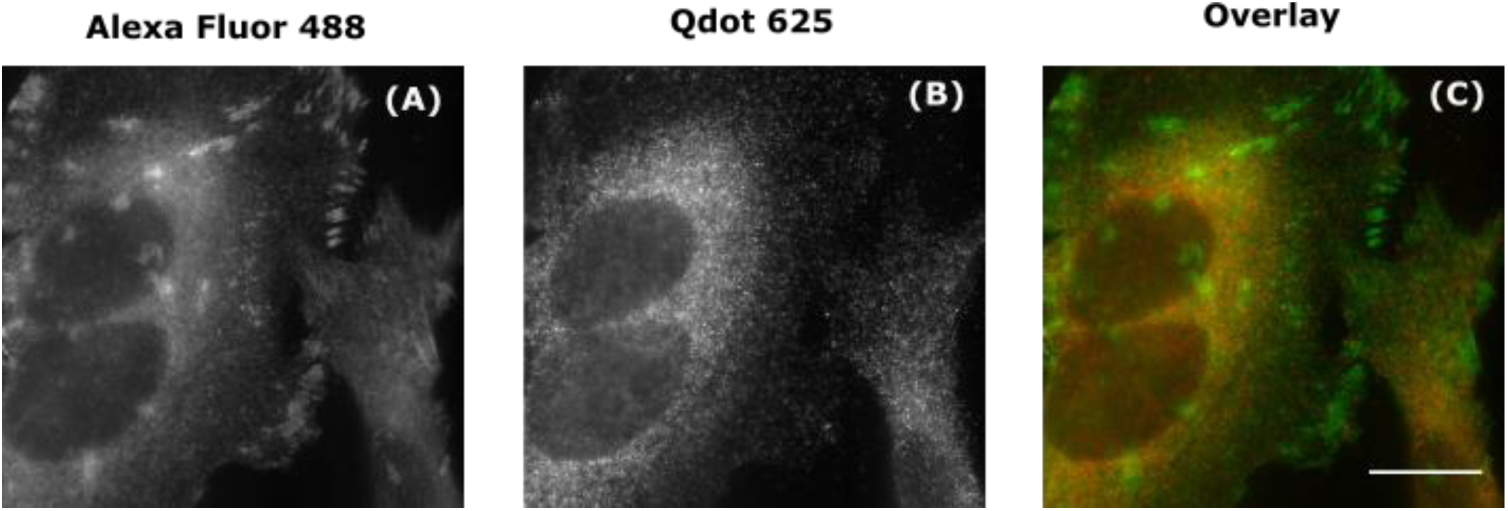
Non-specific labelling of talin with Qdots. Fixed HeLa cells were dual labelled with green Alexa Fluor 488 (A) and red Qdot 625 (B) to produce an overlaid wide-field image (C). Scale bar is 20 μm.

Unlike β-tubulin and talin, SC35 is not only intracellular, but contained within the nucleus, which is crowded with deoxyribonucleic acid (DNA) and proteins. Under conditions identical to the previous experiments, the labelling of SC35 with the Qdot-Ab was non-specific, diffuse, and predominately cytosolic (Figure 4). The intensity correlation of Qdot 625 and Alexa Fluor 488 labelling of SC35 was assessed by plotting fluorescence intensities (Figure 4D) within the overlaid image (Figure 4C). A Pearson’s Correlation Coefficient (r²) value of 0.006 suggested that there was practically no correlation between Qdot 625 and Alexa Fluor 488 signals. Once more, the degree of co-localisation was quantified using the Manders’ coefficient. The average Manders’ coefficient of Alexa Fluor 488 overlapping with Qdot 625 (M1) was 0.19 (SD=0.035, N=3) and the average Manders’ coefficient of Qdot 625 overlapping with Alexa Fluor 488 (M2) was 0.08 (SD=0.031, N=3), which confirms that the labelling of SC35 with Qdot 625 showed no co-localisation with Alexa Fluor 488. To control for unspecific binding of the Qdot-Ab, a negative control was introduced, whereby the cells were incubated with the Qdot-Ab only. In the absence of a primary antibody, there was negligible labelling detected with the Qdot-Ab, which suggests that the SC35 cytosolic signal is due to the presence of minority pool of SC35 in the cytosol.

**Figure 4.**
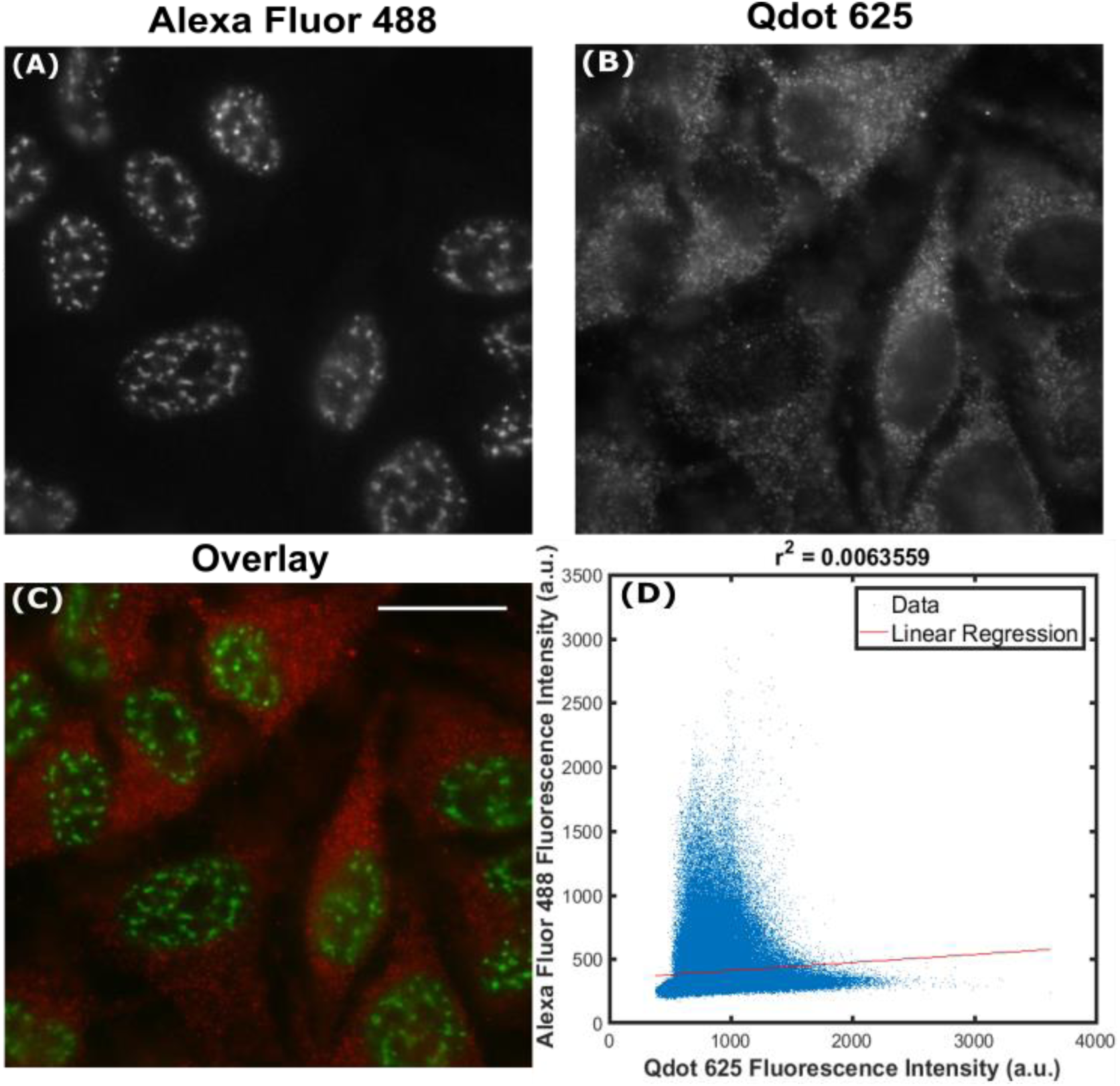
Non-specific labelling of SC35 with Qdots. Fixed HeLa cells were dual labelled with green Alexa Fluor 488 (A) and red Qdot 625 (B) to produce an overlaid wide-field image (C). Scale bar is 20 μm. As a measure of co-localisation between Alexa Fluor 488 and Qdot 625, fluorescence intensities of the overlaid wide-field image (C) were analysed using a custom-written Matlab code to produce a correlation scatter plot (D). Pearson’s Correlation Coefficient (r^2^) of 0.006 indicates no co-localisation. The average Manders’ coefficient of Alexa Fluor 488 overlapping with Qdot 625 (M1) was 0.19 (SD=0.035, N=3) and the average Manders’ coefficient of Qdot 625 overlapping with Alexa Fluor 488 (M2) was 0.08 (SD=0.031, N=3), as determined using JACoP.

*Optimising sample preparation*: We suspected that the size of the commercial Qdot-Ab and the associated accessibility to targets was the reason behind the inability to label complex cytosolic and nuclear structures. Since Qdot-Abs may be sensitive to certain fixation protocols [13, 19, 30], as well as using paraformaldehyde, ice cold methanol was also tried. Methanol fixation and harsher permeabilisation with up to 1% Triton X-100 was used in an attempt to increase accessibility of the Qdots to complex intracellular antigens. Although methanol fixation gave better signal-to-noise ratio and specific Qdot 625 labelling of β-tubulin (Figure 2B), without the need for further permeabilisation, there was only non-specific labelling observed for talin and SC35.

*Labelling with an alternative Qdot 625:* As an alternative of smaller dimensions than the secondary Ab, Qdot 625 conjugated to the biotin-binding protein streptavidin (Qdot-Streptavidin) (Thermo Fisher Scientific, UK) was evaluated. To test the specificity of Qdot-Streptavidin for nuclear targets, a transcription factor, which localises in the nucleus as speckles (sub-nuclear foci), known as hypoxia inducible factor two alpha (HIF2α) was labelled. Fixed HeLa cells were transfected with HIF2α tagged with the fusion protein EGFP (EGFP-HIF2α), incubated with anti-GFP biotin primary antibody, and Qdot-Streptavidin. All of the endogenous biotin sites in the cell were blocked before addition of the biotinylated anti-GFP primary antibody with an endogenous biotin-blocking kit (Thermo Fisher Scientific, UK). Similar results were obtained as previously for Qdot-Ab, with Qdot-streptavidin binding to any cytosolic pool of HIF2α without labelling the distinct speckles in the nucleus (Figure 5). Since the transfected cells have a cytosolic pool of EGFP-HIF2α, this was labelled by Qdot-streptavidin more than in the untransfected cells, where there was no GFP present; hence the very bright fluorescent Qdot signal in the cytosol.

**Figure 5.**
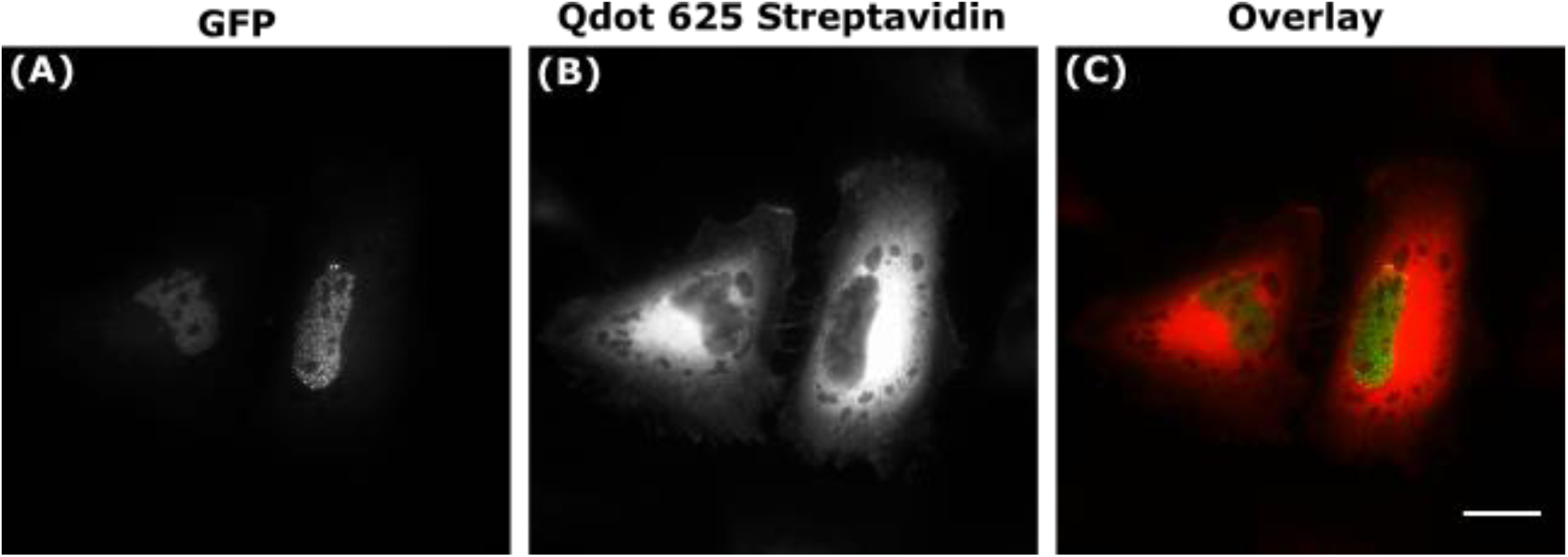
Non-specific labelling of HIF2α with Qdot-Streptavidin. Fixed HeLa cells were transfected with EGFP-HIF2α (A), incubated with a primary anti-GFP biotinylated antibody and Qdot 625 streptavidin conjugate (Qdot-Streptavidin) (B), to produce an overlaid image (C). Scale bar is 20 µm.

*Assessing nuclear accessibility of Qdots*: Interestingly, in all of the Qdot 625 images, even when the staining was non-specific, there was no labelling within the nucleus (Figures 1-5). This suggested that, specificity issues aside, the Qdot-Abs could not access the nucleus at all. This hypothesis was tested by transfecting cells with an unconjugated soluble GFP, which diffuses throughout the cytoplasm and nucleus. We then labelled with either a direct anti-GFP Qdot 625 conjugate (Qdot-Ab) or indirectly with a primary anti-GFP antibody and Qdot-Ab. Both approaches yielded homogenous labelling in the cytosol, with the Qdot signal being excluded from the nucleus (Figure 6). There was also little non-specific Qdot 625 staining in the non-transfected cells. Unlike Qdot 625, when the unconjugated soluble GFP was immunolabelled with a secondary antibody coupled to the fluorescent dye cyanine 3 (Cy3), there was labelling in both the cytosol and nucleus (Figure 6E). To assess the extent at which Qdot 625 did not label soluble GFP within the nucleus, fluorescence intensities of Qdot 625 and GFP were plotted from line scans taken across a section of the cell, including the cytosolic region and nucleus (Figure 6K). There was an obvious decline in the fluorescence intensity of Qdot 625 in the nucleus. Under continuous illumination, the fluorescence intensity of Qdot 625 in the cytosol was greater than that of GFP, due to the superior brightness of Qdots. Multichannel images were taken without adjustment to the focus to rule out different focal planes explaining the absence of Qdots from the nucleus.

**Figure 6.**
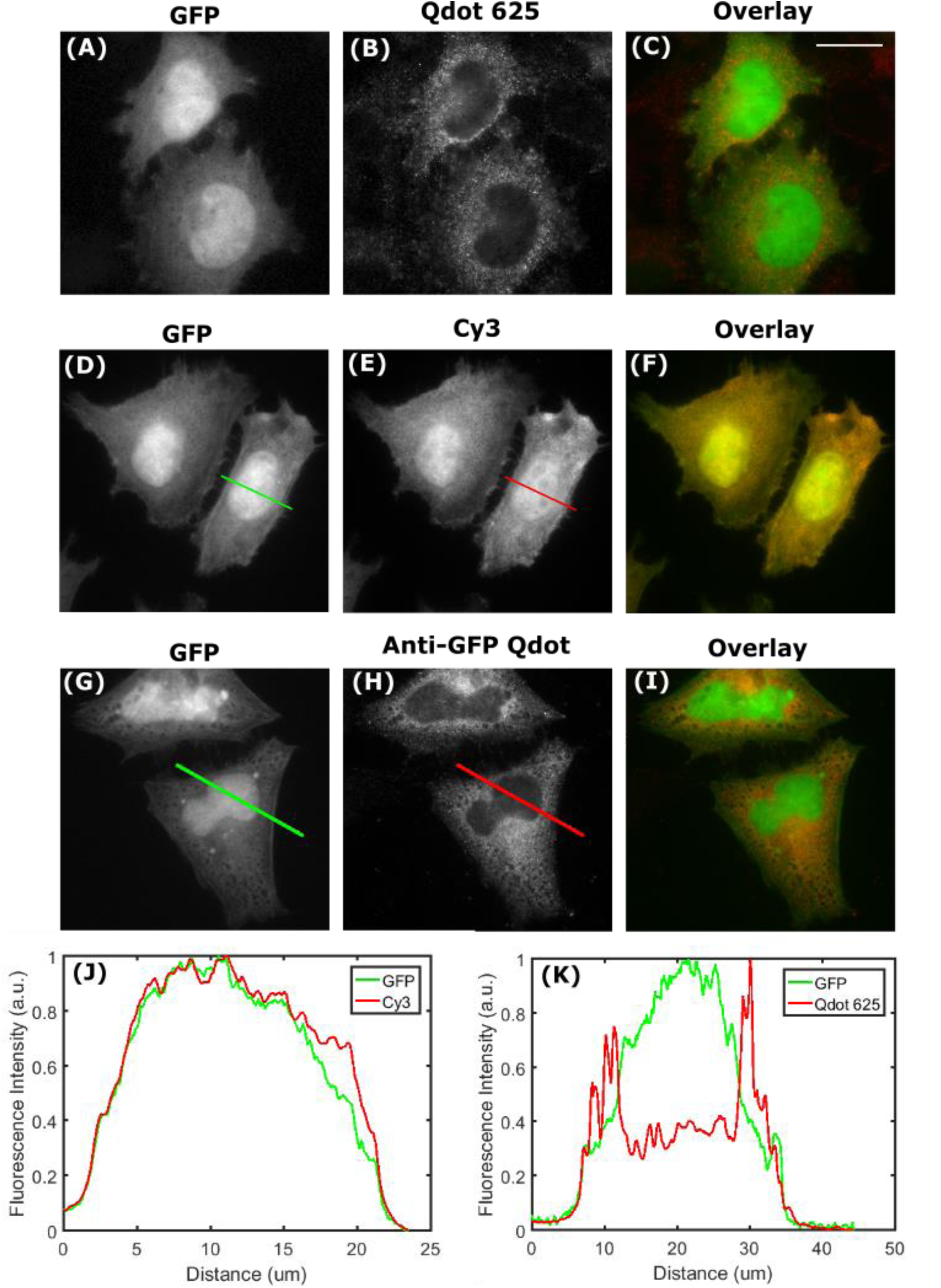
Qdot-Abs are unable to access the cell nucleus. Fixed HeLa cells were transfected with unconjugated soluble GFP (A), incubated with a primary anti-GFP antibody, and red Qdot-Ab (B) to produce an overlaid image (C). Fixed HeLa cells were also transfected with unconjugated soluble GFP (D), incubated with a primary anti-GFP antibody, and red cyanine 3 conjugated to a secondary antibody (E) to produce an overlaid image (F). Direct ICC was done by incubating fixed HeLa cells, transfected with unconjugated soluble GFP (G), with an anti-GFP Qdot 625 conjugate (H) to produce an overlaid image (I). Normalised fluorescence intensities of unconjugated soluble GFP labelled with Cy3 (J) and Qdot 625 (K) were plotted from corresponding lines scans to show no labelling within the nucleus with Qdot 625. Scale bar is 20 μm.

## Conclusion

Fluorescent Qdots are bright and photostable; hence have been advertised as having a lot of potential for imaging in biology, including for the immunolabelling of cellular structures. Yet, several years after promising articles and reviews, Qdots have not found routine use in biological research. Here, we examine the performance of some of the few Qdot conjugates commercially available for immunofluorescence. While all tested antigens could be labelled using antibodies conjugated to fluorescent dyes (Dye-Abs), such as Alexa Fluor 488, only tubulin and extracellular fibronectin could be labelled specifically with Qdot-Abs. Neither a nuclear protein (SC35), nor an antigen within a large complex (talin) could be labelled with the Qdot-Abs. Our current explanation for these results is that the specificity of Qdot-Abs is dependent upon the type of protein and its location within the cell. This can be explained based on the size of the Qdot-Abs and thus steric hindrance leading to limited access to certain epitopes. Additionally, if cross-linking of the target protein has occurred, Qdot 625 aggregates may restrict access to the epitope, and thus affect the specific labelling of these proteins [31]. We conclude therefore, that these Qdot-Abs are not suitable to detect complex intracellular structures, unless the proteins are abundant and have multiple accessible antigens along the structure; although this contradicts previous literature by Montόn *et al.,* (2009), who found that Qdot-Abs were more specific for proteins that are scarce in the cell [20]. Overall, the specific labelling of certain proteins, such as SC35 and talin, with commercial Qdot-Abs was reproducibly non-existant.

Beyond those commercially available QDs, there have been a number of ICC protocols published, for the labelling of different proteins with Qdot-Abs [6, 20, 24], but they have usually focused on the unproblematic labelling of tubulin for their proof of principle experiments. We suggest that future developments of Qdot-Abs include other more challenging targets as benchmarks, e.g. those evaluated in this article. The ideal scenario would be a toolbox of commercially available Qdot-Abs that can consistently be used to label any biological structure of interest and not just those on the apical surface of cells, much in the same way as Dye-Abs are used.

## Methods

### Cell Culture

Human cervix epitheloid carcinoma (HeLa, ECACC number 930210a3) cells were cultured in a 75 cm^2^ flask at 37 °C with 5 % CO2, minimum essential media (MEM, Life Technologies, UK) supplemented with 10% (v/v) foetal calf serum (FCS), and 1% non-essential amino acids (NEAA). Cells were split 1,000,000 cells/mL when ≥80% confluent with trypsin-EDTA. Rat mammary (Rama) 27 fibroblasts were cultured in a 75 cm^2^ flask at 37 °C with 5% CO2, Dulbecco’s modified Eagle’s medium (DMEM, Life Technologies, UK) supplemented with 10% (v/v) FCS (Life Technologies, UK), 0.75% (w/v) sodium biocarbonate, 4 mM L-glutamine, 50 ng/mL insulin, and 50 ng/mL hydrocortisone (Sigma-Aldrich, UK), as described previously [32]. Cells were split 1:8 when ≥60% confluent with trypsin-EDTA. A stable cell line TC7 3xGFP (expressing tubulin-GFP) was cultured in a 75 cm² flask at 37 °C with 5% CO2, MEM (Life Technologies, UK) supplemented with 10% (v/v) FCS, 1% NEAA, and genetitin (Sigma-Aldrich, UK), as described previously [33]. Cells were split 1:15 when ≥80% confluent with trypsin-EDTA.

### Transfection

HeLa cells were seeded onto 35 mm glass bottom dishes (100,000 cells/mL) and transfected with pG-EGFP-A (soluble GFP) or pG-EGFP-HIF2α (EGFP-HIF2α) using FuGENE6 transfection reagent (Roche Limited, UK), following the manufacturer’s protocol (3:1 transfection reagent:DNA plasmid).

### Site Click Conjugation of Qdot625 to anti-GFP

Following the manufacturer’s protocol, a commercial site-click Qdot 625 antibody conjugation kit (Thermo Fisher Scientific, UK) was used to conjugate a primary mouse (clones 7.1 and 13.1) anti-GFP antibody (Roche Limited, UK) to dibenzocyclooctyne (DIBO) modified Qdot 625. The concentration of the Qdots in the conjugate was calculated to be 3 μM using the equation 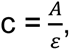 where c is the concentration of DIBO-modified Qdot 625 attached to the primary antibody, A is the absorbance of Qdot 625, and Ɛ is the extinction coefficient of Qdot 625 (500,000 M^-1^cm^-1^). The absorbance between 605 nm-612 nm (step 10 nm) was measured to be 1.5 a.u. using a quartz cuvette with a 1 cm path length, on a SpectraMax 34 Plus spectrophotometer (Molecular Devices, UK).

### Immunofluorescence

An overview of the immunofluorescence procedure is shown in scheme 2. Briefly, cells were seeded onto glass coverslips and grown until confluent, washed once in phosphate buffered saline (PBS) (37 °C), and fixed in 4% (w/v) paraformaldehyde (PFA) for 10 min or 100% ice cold methanol (5 min). Cells were washed 3x in PBS (5 min), permeabilised with 0.25% Triton X-100 in PBS for 60 min (except methanol fixation), washed again 3x in PBS (5 min), and incubated with 6% bovine serum albumin (BSA, Sigma-Aldrich, UK) in PBS (60 min). Primary antibodies produced in mouse (anti-β-tubulin TUB 2.1, Sigma-Aldrich, UK; anti-GFP Roche Limited, UK; anti-SC35, Abcam, UK; and anti-talin, Sigma-Aldrich, UK), those produced in rabbit (anti-fibronectin, Sigma-Aldrich, UK), and biotinylated anti-GFP (Abcam, UK) were diluted 1:100 in 6% BSA and incubated overnight at 4 °C. Cells were washed 3x in PBS (5 min) and incubated simultaneously with either a mixture of F(ab’)2-Donkey anti-mouse IgG H+L secondary antibody Qdot 625 conjugate, Qdot 625 anti-GFP conjugate, F(ab’)2-Goat anti-rabbit IgG H+L secondary antibody Qdot 625 conjugate, or Qdot 625 streptavidin conjugate (Thermo Fisher Scientific, UK), diluted 1:50 to 20 nM in 6% BSA, and goat anti-rabbit Alexa Fluor 488 or goat anti-mouse Alexa Fluor 488 (Thermo Fisher Scientific, UK), diluted 1:500 in 6% BSA; or anti-mouse cyanine 3 (Sigma-Aldrich, UK) diluted 1:500 in 6% BSA, at room temperature (60 min). Before preparation of the Qdot 625 conjugated secondary antibody, the vial was centrifuged at 5,000 g for 3 min to remove any aggregates. After 3 washes in PBS (10 min), coverslips were mounted onto slides with Dako fluorescent mounting media (Dako, UK), and stored at 4 °C. A negative control of Qdot 625 conjugated antibody only was also prepared to show any background staining.

**Scheme 2.**
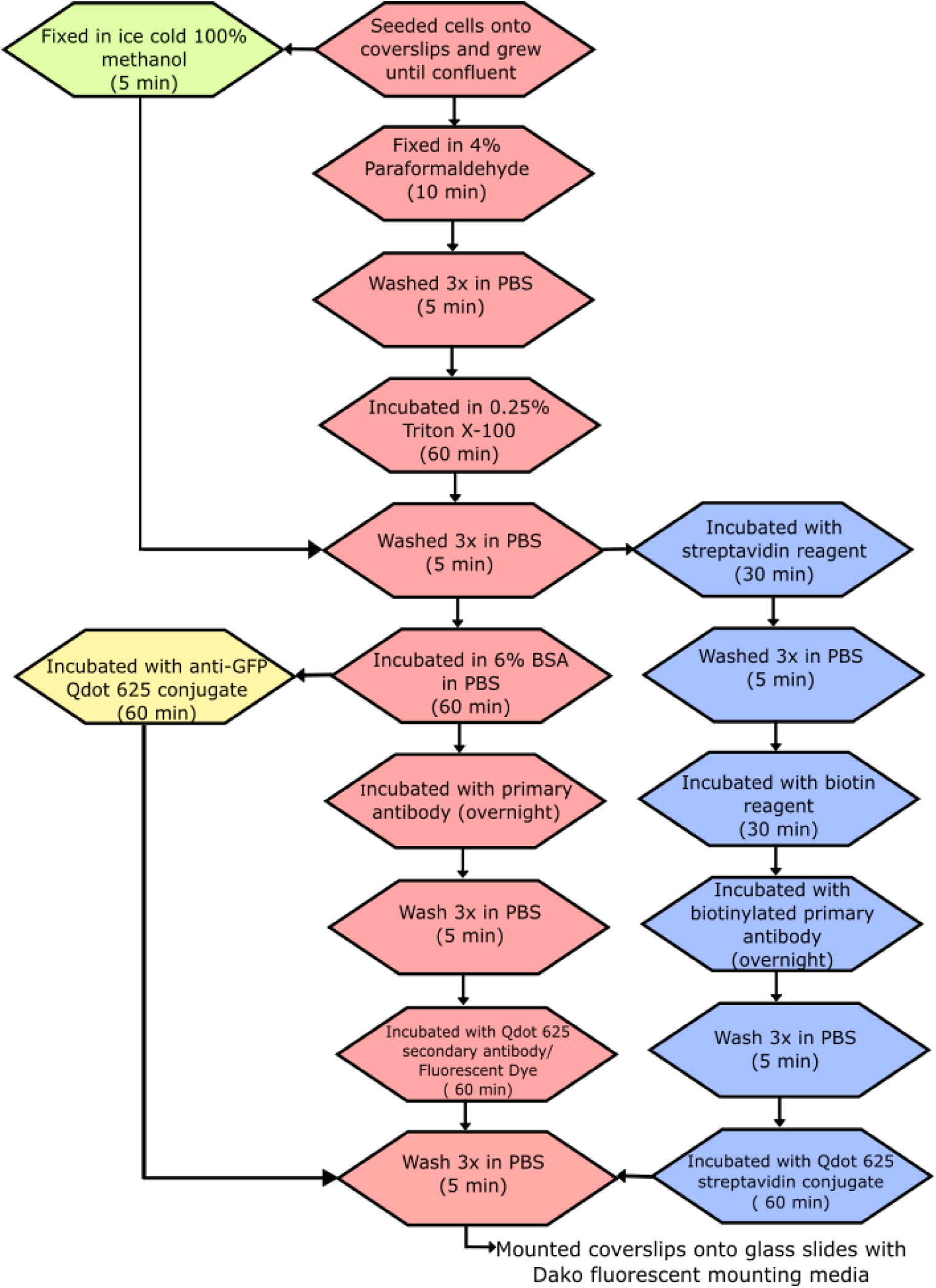
Immunofluorescence Protocol. The pink boxes show the method for use with a primary antibody and Qdot 625/Fluorescent dye conjugated to a secondary antibody, green box is for methanol fixation, yellow boxes for the anti-GFP Qdot 625 conjugate, and blue boxes are for biotinylated primary antibody with Qdot 625 streptavidin conjugate.

### Wide-field Imaging

Images were taken on a wide-field epifluorescence microscope (Carl Zeiss Axio Observer Z.1, Germany) with a 16 μm 512 x 512 pixel sensitive electron-multiplying charge-coupled device (EMCCD) camera (Andor iXon 897 Ultra), polychromatic mercury arc lamp, 39106-AT-QDot 625 filter set (Chroma Technology Corporation, USA), and a 100x 1.45 NA oil-immersion objective. Fluorescent and corresponding brightfield images were acquired using Micro-Manager software [34]. The same acquisition settings were used for each set of images, including lamp power, exposure time, and gain.

### Co-localisation Analysis

Pearson Correlation Coefficient scatter plots of Qdot 625 and Alexa Fluor 488/GFP were produced using a custom-made Matlab code [35]. Manders’ Correlation Coefficients were determined using a Just Another Co-localization Plugin (JACoP) [29] in FIJI [36].

## Acknowledgements

We express gratitude to Professor Chöle Bulinkski from Colombia University for the TC7 3xGFP cells and also appreciation to Professor Philip Rudland from University of Liverpool for supplying the rat mammary (Rama) 27 fibroblasts. We would like to acknowledge and thank the Liverpool Centre for Cell Imaging (CCI) for training and access to equipment. The Epifluorescence microscope used in this study was funded in part and DM was fully funded by the Medical Research Council (Grant number: MR/K015931/1). All of the experiments were carried out using funding from an iCASE BBSRC studentship.

